# Neural responses to happy, fearful and angry faces of varying identities in 5- and 7-month-old infants

**DOI:** 10.1101/2020.04.07.030395

**Authors:** Laurie Bayet, Katherine L. Perdue, Hannah F. Behrendt, John E. Richards, Alissa Westerlund, Julia K. Cataldo, Charles A. Nelson

## Abstract

Facial emotion processing is an important social skill that develops throughout infancy and early childhood. Here we investigate the neural underpinnings of the ability to process facial emotion across changes in facial identity in cross-sectional groups of 5- and 7-month-old infants. We simultaneously measured neural metabolic, behavioral, and autonomic responses to happy, fearful, and angry faces of different female models using functional near-infrared spectroscopy (fNIRS), eye-tracking, and heart rate measures. We observed significant neural activation to these facial emotions in a distributed set of frontal and temporal brain regions, and longer looking to the mouth region of angry faces compared to happy and fearful faces. No differences in looking behavior or neural activations were observed between 5- and 7-month-olds, although several exploratory, age-independent associations between neural activations and looking behavior were noted. Overall, these findings suggest more developmental stability than previously thought in responses to emotional facial expressions of varying identities between 5- and 7-months of age.

---

The discrimination and identification of facial expressions of emotion form an important channel for human non-verbal communication. The behavioral ability to differentiate among different facial expressions (such as between happy and angry or fearful faces) is thought to emerge between 5 and 7 months in typically developing infants, a potential first step in the developmental emergence of facial expression understanding (Leppänen & Nelson, 2009). For example, 7-month-olds differentiate between some expressions (e.g. happy versus fear, happy versus angry, or fear versus angry) in behavioral paradigms, even when expressions are modelled by faces of different identities (Bayet & Nelson, 2019; Leppänen & Nelson, 2009).

Eye-tracking studies suggest that infants’ developing abilities to differentiate between facial expressions mature along with their ability to focus on internal facial features, such as the eyes and mouth (Hunnius, de Wit, Vrins, & von Hofsten, 2011; Soussignan et al., 2017), that are central to human communication (Gliga & Csibra, 2007) and convey distinguishing information about facial expressions (Smith, Cottrell, Gosselin, & Schyns, 2005). Preliminary evidence additionally suggests that individual differences in eye-looking durations (e.g. infants being eye or mouthlookers) relate to differences in facial emotion discrimination abilities (Amso, Fitzgerald, Davidow, Gilhooly, & Tottenham, 2010). Looking behavior, and in particular looking towards the eyes of emotional faces, may thus drive or reflect infants’ abilities to extract information from and discriminate between facial emotions (Bayet & Nelson, 2019).

Heart rate has been found to index attention in infancy (Courage, Reynolds, & Richards, 2006; Richards & Casey, 1991), including attention to facial emotions. Transient heart rate decelerations are thought to reflect attentional orienting (Perdue, Edwards, Tager-Flusberg, & Nelson, 2017), with larger and longer heart rate deceleration to fearful as compared to non-fearful faces by 7 months of age (Peltola, Hietanen, Forssman, & Leppänen, 2013).

Functional near-infrared spectroscopy (fNIRS) has provided some information about which cortical regions of the infant brain are specifically activated in response to facial expressions of emotion. In adults, processing facial expressions engages a distributed set of coactive brain regions (Fusar-Poli et al., 2009; Kesler-West et al., 2001) that include the fusiform face area, superior temporal gyrus (STG; Narumoto, Okada, Sadato, Fukui, & Yonekura, 2001; Winston, Henson, Fine-Goulden, & Dolan, 2004), amygdala, and frontal cortices (Nakamura et al., 1999; Vuilleumier, Armony, Driver, & Dolan, 2001). While not all brain regions implicated in emotional face processing in adults are accessible by fNIRS due to limited depth penetration, superficial temporal and frontal cortices can be accessed. Prior infant fNIRS work has shown differential brain responses to happy and angry faces in the temporal cortex from 6-7 months of age (Nakato, Otsuka, Kanazawa, Yamaguchi, & Kakigi, 2011) and differential responses to happy, angry, and/or fearful faces in the right inferior frontal cortex at 7-months of age when modelled in conjunction with certain individual epigenetic differences (Grossmann, Missana, & Krol, 2018; Krol, Puglia, Morris, Connelly, & Grossmann, 2019). fNIRS has also revealed that medial frontal regions are implicated in processing happy faces in infants from 9-13 months (Minagawa-Kawai et al., 2009) and linked individual differences in frontal responses to emotional faces with earlier epigenetic changes (Krol et al., 2019) and later behavior (Grossmann et al., 2018). However, no studies to date have directly tested for differences in neural activity between 5 and 7 months, a critical time window during which differential responses to fearful faces are often first observed with ERP (Leppänen, Richmond, Vogel-Farley, Moulson, & Nelson, 2009; Xie, McCormick, Westerlund, Bowman, & Nelson, 2018). In addition, no prior infant fNIRS study has examined responses to facial emotions when facial identity varies, a focus of classic behavioral studies in this age group (Bayet & Nelson, 2019; Leppänen & Nelson, 2009), or in relation to simultaneous looking behavior to facial features. Such a test would greatly extend past work by providing a more robust indicator of facial emotion discrimination than responses to facial expressions of emotions from the same model. The current study sought to address these gaps by examining the neural, behavioral, and autonomic correlates of emotional face processing in cross-sectional groups of 5- and 7-month-old infants. In line with existing studies in this field, we focused on canonical happy, fearful and angry facial expressions. Facial identity varied within each trial to isolate responses to emotional categories independently of the identity of the model, rather than to expression changes of a single model.

## Materials and Methods

### Participants

Cross-sectional groups of 48 5-month olds (26 males, mean age 153, ± 4 days) and 52 7-month olds (30 males, mean age 212 ± 5 days) formed the final sample. All were typically developing, with no known pre- or perinatal complications, and born after 37-weeks of gestation. Data from an additional 23 5-month-olds and 34 7-month-olds were collected but excluded due to cap refusal (n=2 5-month-olds, n=1 7-month-old), completing fewer than 3 trials per condition (n=3 5-month-olds), inaccurate cap placement (deviation from ideal by more than 1.5 cm in any direction as determined by post-hoc review of placement photos; n=5 5-month-olds, n=10 7-month-olds), over 25% (12 of 46) channels unusable (n=11 5-month-olds, n=19 7-month-olds), technical malfunction (n=1 5-month-old, n=4 7-month-olds) or experimenter error (n=1 5-month-old). Additionally, 5 participants were excluded due to self-report of maternal opioid or antipsychotic medication use during pregnancy (n=3 5-month-olds) or subsequent ASD diagnosis (n=1 5-month-old, n=2 7-month-old). Eye-tracking data from 2 included participants were excluded due to inability to calibration failure (n=1 7-month-old) or technical problems with the eye-tracker (n=1 7-month-old). The study was approved by the Institutional Review Board of Boston Children’s Hospital. Written informed consent was provided by the infant’s parent before starting the session.

### Stimuli

Photographs of women with happy, fearful and angry face expressions from the NimStim dataset (Tottenham et al., 2009) were presented in a block design. Faces were in full color with open mouths and visible teeth. In each block, 5 faces of different women with the same expression were presented for 1-second each with a random inter-stimulus interval of 0.2-0.4 seconds. The race of the women in the presented stimuli was matched to maternal self-reported race. Stimulus blocks were followed by a 10-second full color video of abstract moving shapes. Up to 10 blocks were presented for each emotional condition for a total of up to 30 blocks as tolerated.

### Experimental Procedure

fNIRS data were collected using a Hitachi ETG-4000 continuous-wave system (wavelengths at 695 nm and 830 nm) with 46 channels distributed over the bilateral frontal and temporal regions (**Figure 1**; source-detector distances approximately 3 cm), sampling at 10 Hz. Simultaneous binocular eye-tracking data was recorded with a Tobii T120 eye-tracker sampling at 60Hz.

**Figure 1:**
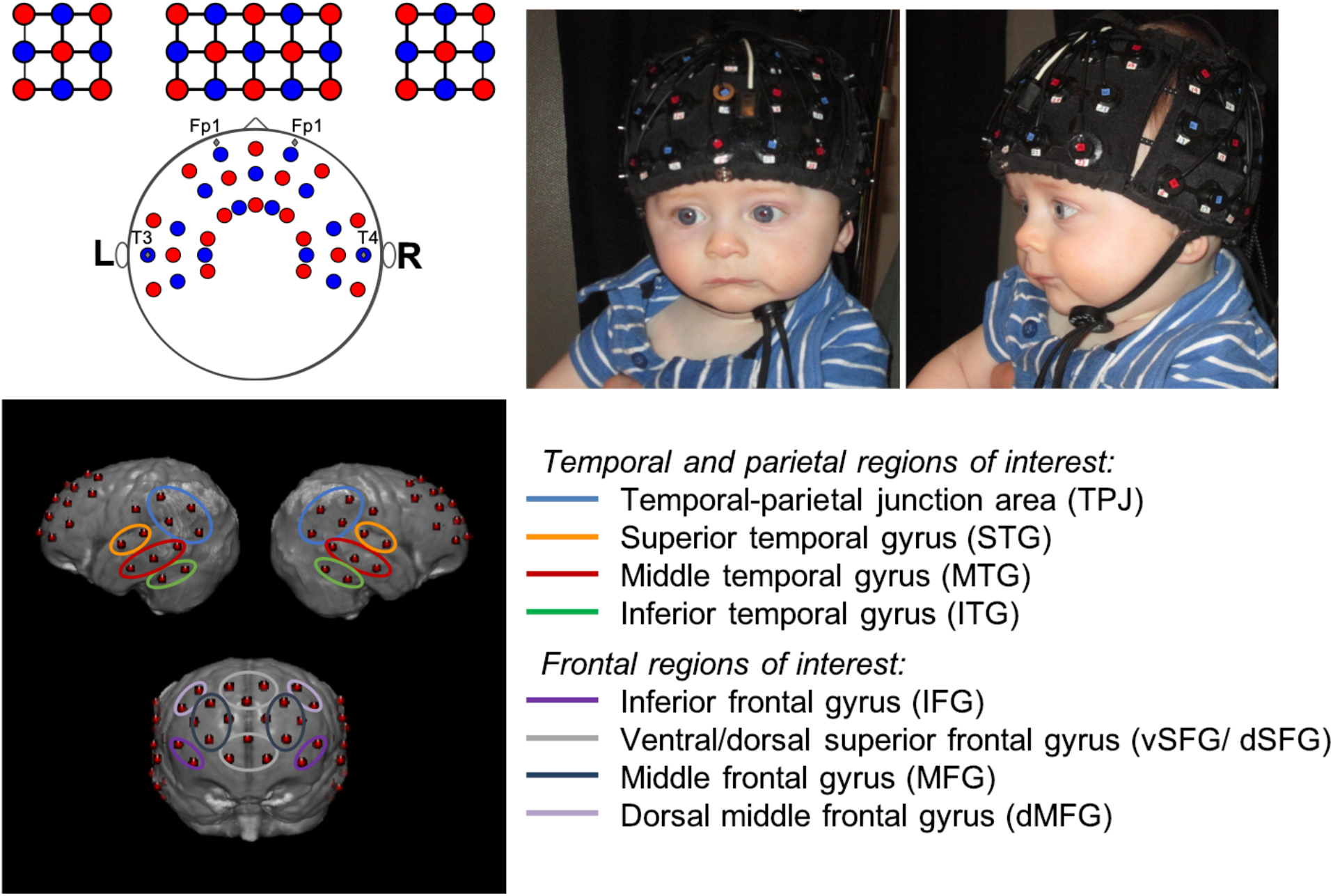
Top left, diagram of sources (red) and detectors (blue) and their placement relative to the 10-20 system. Top right, front and side photo of probe on 7-month-old infant. Bottom, modeled channel locations and ROI designations displayed on a 7.5-month-old MRI atlas (Richards et al. 2016).

Infants were held on a caregiver’s lap while they viewed a 17-inch screen approximately 60 cm away. Face stimuli were presented at a visual angle of 14.3° high by 12.2° wide. Parents wore visors to block their view of the presented stimuli and were asked to refrain from talking to the infants during the experiment.

Eye-tracker calibration was performed using a 5-point calibration procedure. Once sufficient eyetracking calibration was obtained (minimum of 3 out of 5 locations) or failed after two attempts (in which case the eye-tracking data was excluded, n = 1 7-month-old), the paradigm began. Stimulus presentation was experimenter-controlled in an adjacent room using E-Prime 2.0 (Psychological Software Products, Harrisburg, PA). Blocks were initiated when the infant was looking at the screen. Another experimenter was seated next to the infant, redirecting attention to the screen between blocks if necessary.

### Video coding

Infants were video recorded during the experiment. Videos were coded offline for percent looking to the screen by trained coders. Infants were considered to have attended to a trial block if they looked at the screen for at least 2.5 seconds of the presented face stimuli (5 seconds). Inter-coder agreement was assessed from a randomly selected 20% of the sample. Average inter-coder agreement was 95% for trial include/exclude decision; 80% of trials were within 10% agreement for percent looking between coders. Percent looking to the screen as determined by video coding was only used for fNIRS data quality control. Baseline periods were not coded for infant attention. However, blocks were only initiated when infants were looking at the screen, so that in the time period immediately preceding the stimulus the infants were looking at the crosshair on the screen. The mean number of accepted blocks per emotional category was 7.1 at 5 months and 7.8 at 7 months. The effect of condition (within-subject factor of emotional category) on the number of accepted blocks, as a main effect or in interaction with age, was not significant (*ps* >.4). On average, 7-month-olds had more accepted blocks (main effect of age, *F*(1, 98)=4.16, *p*=.044) Average video-coded percent looking in accepted trials was 88.3% at 5-months and 88.2% at 7-months. There were no significant effects of condition, age, or their interaction (all *ps* >.2) on percent looking in accepted trials.

Despite good eye-tracking calibration before the experiment was started, some participants had large discrepancies between eye-tracker reported looking and video-coded looking time, usually due to participant position changes during the experiment leading to the calibration no longer being valid. Participants’ data were removed from eye-tracking analyses if they did not have at least 3 blocks for every condition with total looking as measured by eye-tracker in agreement (within 25%) of video coded looking time (n=17 5-month-olds and n=4 7-month-olds), leaving n=30 5-month-olds and n=47 7-month-olds in combined eye-tracking and fNIRS analyses.

### Anatomical localization

Age-appropriate MRIs were chosen from a database (Richards, Sanchez, Phillips-Meek, & Xie, 2016; Sanchez, Richards, & Almli, 2012) to model the underlying anatomy of each channel. A group of 13 MRIs (resp. 15) was used for 5-month-olds (resp. 7-month-olds), with age and head circumference matched to the experimental group (Age of MRI participants: 5 mo 148 ± 13 days, 7 mo 212 ± 17 days, Head circumference of MRI participants: 5 mo 42.8 cm ± 1.6 cm, 7 mo 44.4 ± 2.1 cm). Head probe geometry and cap placement photographs were used to virtually place the optodes on each MRI using previously developed methods (Lloyd-Fox et al., 2014). Photon propagation modeling (Fang, 2010) estimated diffuse optical tomography sensitivity functions from each source-detector pair comprising a channel, confirmed with previously described geometrical methods (Lloyd-Fox et al., 2014; Okamoto & Dan, 2005). Intersecting cortical regions were labeled using the LONI atlas (Fillmore, Richards, Phillips-Meek, Cryer, & Stevens, 2015). Structural regions of interest (ROIs) were defined based on the averaged localization of each channel over the group of MRIs. As localization was similar for the 5- and 7-month-olds, the same ROIs were used across ages. Channel location estimation was also done on an average MRI 7.5 month-old template for visualization (Richards et al., 2016). ROIs contained 2-4 channels each, subdividing the bilateral Inferior (ITG), Middle (MTG), and Superior Temporal Gyri (STG), bilateral Temporal-Parietal Junction (TPJ), bilateral Inferior (IFG), Middle (MFG), and dorsal Middle (dMFG) Frontal Gyri, and medial ventral (vSFG) and dorsal (dSFG) Superior Frontal Gyri (**Figure 1**). Participants had to have greater than 50% of the channels valid in each ROI to be included in the group analysis for a particular region, leading to some non-uniformity in number of participants included in each ROI.

### fNIRS data analysis

fNIRS data analysis was performed using Homer2 (Huppert, Diamond, Franceschini, & Boas, 2009) in MATLAB. Unusable channels were defined as having values greater than 98% or less than 2% of the total raw range of intensity data for more than 5 seconds and excluded. Raw data were converted to optical density, and wavelet motion correction with iqr=0.5 was applied (Behrendt et al. 2018). Residual motion artifacts were identified (tMotion=1, tMask=1, stdevThresh=50, ampThresh=1); affected blocks were excluded. Data were bandpass filtered from 0.05-0.8 Hz to eliminate drift and cardiac artifact. The modified Beer-Lambert law (DPF=5) was applied to convert optical density to chromophore concentration (Duncan et al., 1995). Blocks were extracted from 2-seconds prior to 16-seconds after-stimulus onset, using the pre-stimulus time for baseline correction. As blocks were spaced 18-seconds apart or more, they did not overlap. Concentrations were z-scored by dividing by the standard deviation over the pre-stimulus baseline period.

The grand mean hemodynamic response over all participants and ROIs was used to select a timewindow of interest ± 2 seconds around its peak. Group responses were statistically assessed by one sample *t*-tests vs. zero response, corrected for multiple comparisons at the False Discovery Rate (alpha=0.05).

Block-related heart rate changes were extracted from the fNIRS data using previously described methods (Perdue, Westerlund, McCormick, & Nelson, 2014). Changes in heart rate from the baseline period were calculated by averaging over the first 6-seconds of the trial, when the stimulus was presented. Changes in heart rate could be positive or negative; positive heart rate responses (accelerations) would indicate increases in arousal, while negative heart rate responses (decelerations) would indicate increases in attention.

### Eye-tracking analysis

Rectangular “eyes” and “mouth” areas of interest (AOI) of equal sizes were defined, bounded by the sides of the face and by either the hairline on the top and the midpoint of the face as the bottom (“eye” AOI), or the midpoint of the face on the top and the chin of the face as the bottom (“mouth” AOI). Total looking time to the eye and mouth AOIs while the face stimulus was on the screen was calculated for each block, then averaged over blocks for each condition and participant. In contrast to using percent looking time to the eye and mouth as a percentage of total looking time, this approach was meant to derive an index of the amount of “eye” or “mouth” visual information processed rather than characterize the relative allocation of attention to each feature controlling for total looking time.

### Statistical analysis

Participants were included in a particular analysis if they had complete data to fit the model being tested. Differences in oxyHb responses, looking times, mouth-looking times, and heart rate responses based on the effects of condition (happy, fear, angry; within-subject), age (5-months, 7-months; between-subject), and their interaction were tested using repeated-measures ANOVAs. For oxyHb responses, separate models were run for each ROI. When no significant interaction term was found, the interaction was removed, and the model was rerun. Separate models were run to test for condition-by-eye-looking or condition-by-mouth-looking effects on oxyHb responses in each ROI.

## Results

### fNIRS activations

Grand-average peak oxyHb activation occurred at 8.6 seconds post-onset (**Figure 2**; 8.7 seconds in 5-month-olds, 8.4 seconds in 7-month-olds). Therefore, a time-window of interest from 6.6-10.6 seconds was chosen to extract mean oxyHb activations for each participant in each condition and ROI. To align with prior work in infants, our analysis focused on oxyHb activation, but we also report deoxyHb for completeness. Grand-average peak deoxyHb activation (negative deflection) occurred at 9.5 seconds post-onset (9.5 seconds in 5-month-olds, 9.4 seconds for 7-month-olds).

**Figure 2:**
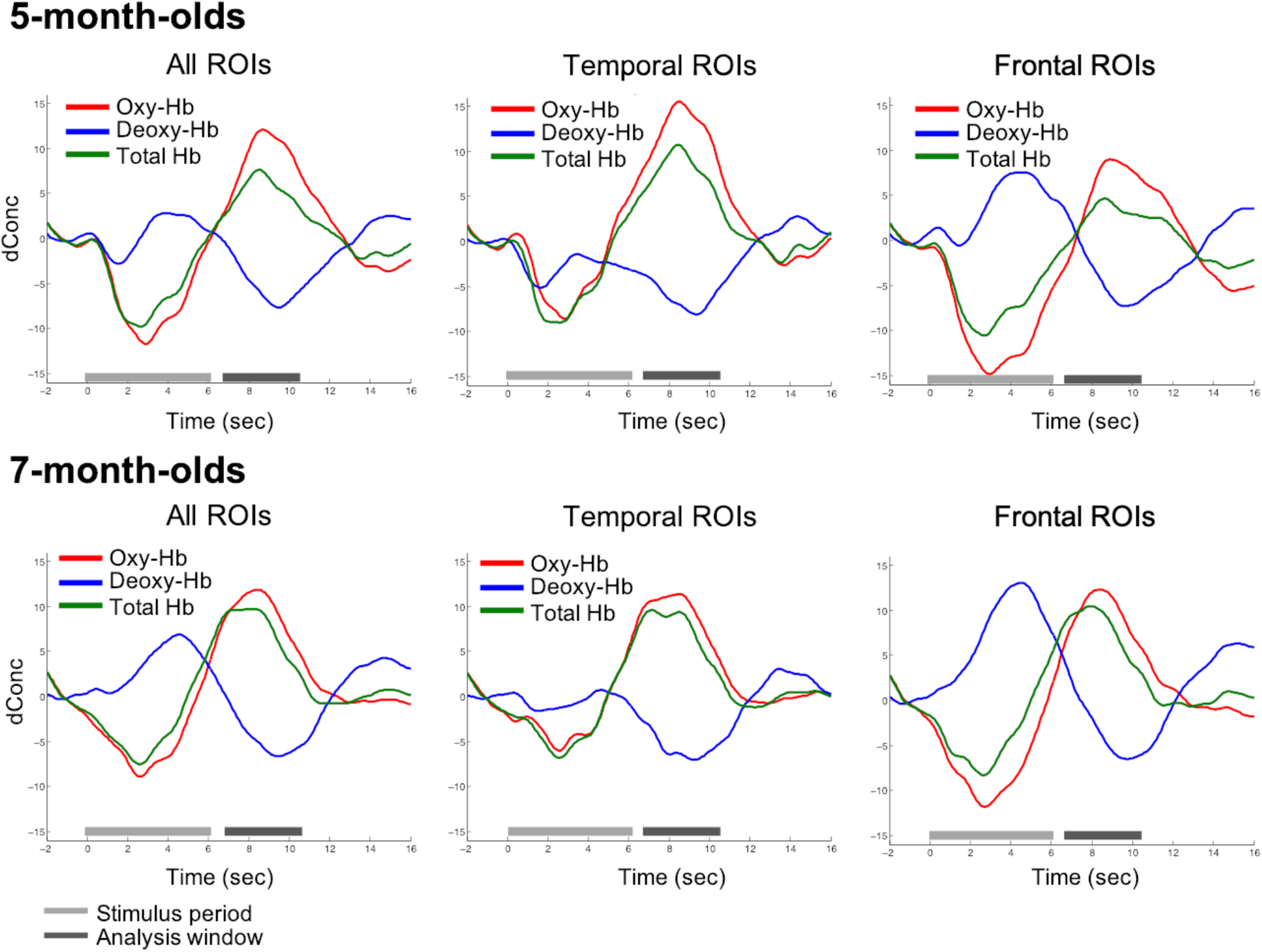
Time-course of grand-mean hemodynamic response averaged over conditions, participants, and ROIs for each age.

Significant activations after FDR correction are reported in **Table 1** and **Figure 3**. Significant oxyHb activation to happy faces was found in the vSFG, dSFG, right dMFG, and bilateral MFG, IFG, MTG, STG, ITG and TPJ. OxyHb activation to fearful faces was seen in the bilateral MTG, rTPJ, and left ITG. OxyHb activation to angry faces was seen in the left MFG and ITG, and in the bilateral IFG and MTG. Significant deoxyHb activations to happy faces were seen in the bilateral MFG, bilateral IFG, left dMFG, left STG, and right TPJ, and in the left MTG for fearful faces. No significant deoxyHb activations were seen to angry faces after FDR correction. Results in each age group are reported in Supplemental Tables 1-2 and Supplemental Figures 1-2.

**Table 1:**
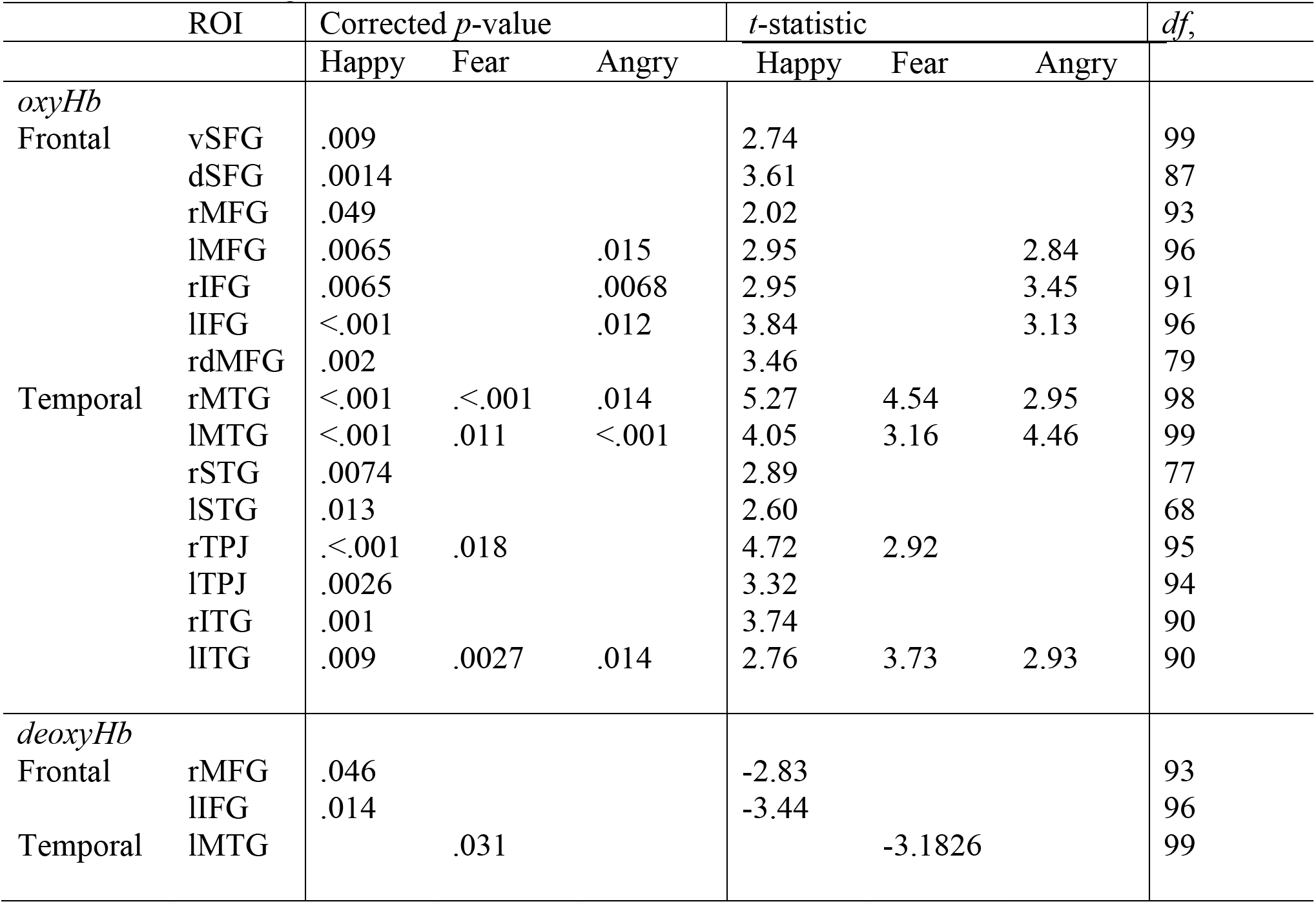
ROIs with significant activation in combined 5- and 7-month-old cohort

**Figure 3:**
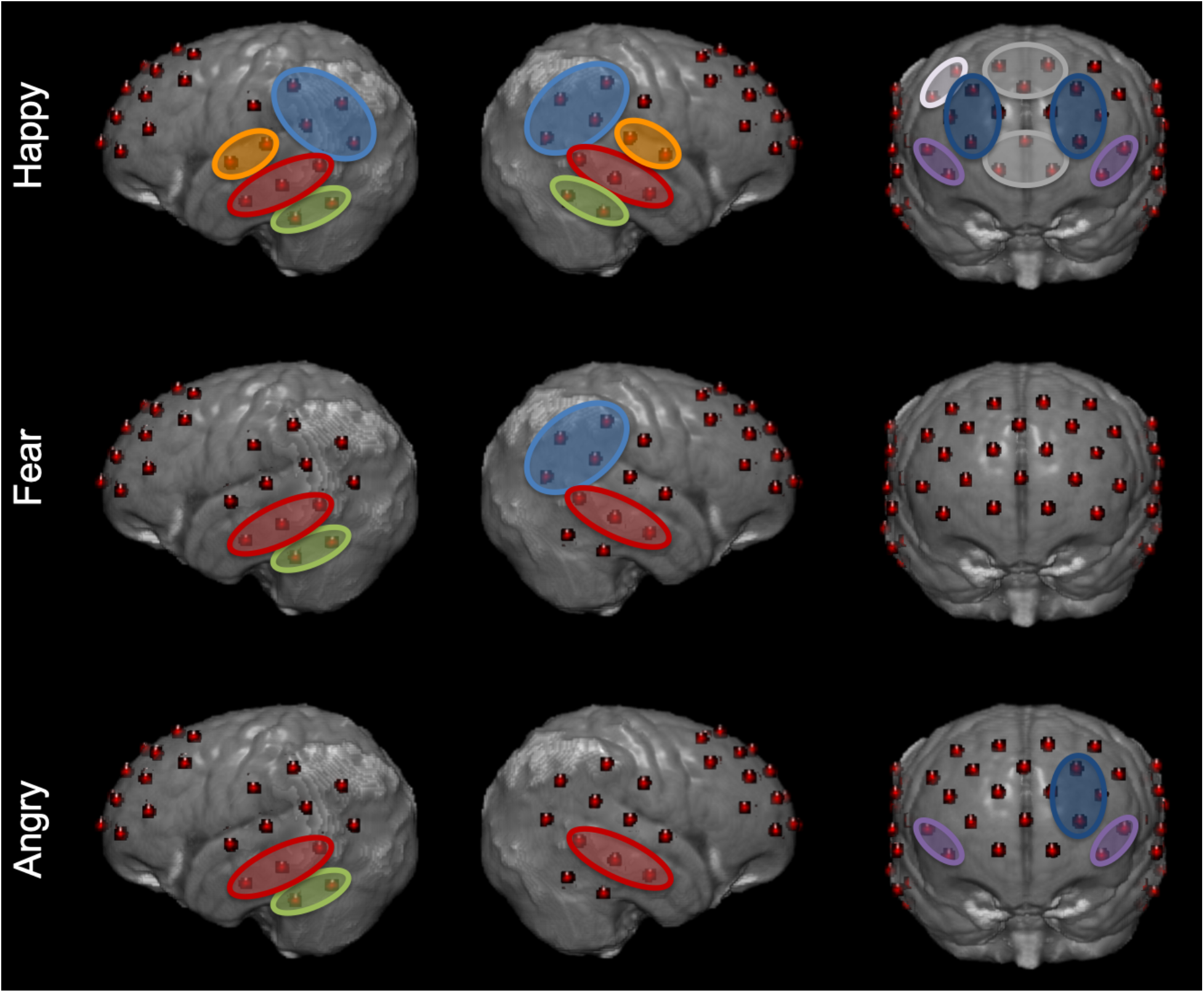
Significant oxyHb activations for each emotional category after correcting for multiple comparisons displayed on a 7.5-month-old MRI atlas (Richards et al. 2016).

No significant effect of age or age-by-condition interactions on oxyHb responses were found in any ROI. There was an exploratory effect of condition on oxyHb responses in the dSFG, with greater activation to happy versus fearful faces (Supplementary Results, Supplementary Figure 3). However, this effect did not survive FDR-correction for multiple comparisons over ROIs.

### Eye-tracking

Infants looked to the eyes for 3.5-seconds and to the mouth for 1.0-seconds on average during included blocks. Eye-looking and mouth-looking were negatively correlated (*r^2^*=.53, *p*<0.001). No significant effect of age or age-by-condition interaction on eye-looking or mouth-looking were found. A marginally significant effect of condition on eye-looking was found (*F*(2, 224)=2.85, *p*=.060), with more looking to the eyes of fear faces compared to angry faces (t(75)=2.33, *p*=.023) but no significant difference in eye-looking between happy and angry (*t*(75)=0.73, *p*=.46) or happy and fear faces (*t*(75)=1.62, *p*=.11). A significant effect of condition on mouth looking was found (*F*(2, 224)=4.40, *p*=.013), with more looking to the mouth of angry faces as compared to happy (*t*(75)=2.39, *p*=.019) and fearful faces (*t*(75)=2.67, *p*=.009), but no difference in mouth-looking between happy and fearful faces (*t*(75)=0.48, *p*=.64). These effects were observed despite all stimuli having open mouths with visible teeth.

### Heart rate responses

No significant effects of age, condition, or their interaction on heart rate responses were found (all *p*s > 0.5)

### Combined fNIRS and eye-tracking analyses

We next investigated whether fNIRS responses varied as a function of the interaction of accumulated looking time to the eyes or mouth and emotion condition, as emotional category affects which feature, eyes or mouth, may be most salient or diagnostic. Differences in oxyHb responses based on looking time were tested using repeated-measures condition-by-accumulated-looking-time ANOVAs in each ROI, separately for eye- and mouth-looking. While we did not find differences in responses due to age in looking behaviors or brain activation on their own, models controlled for age to eliminate the possibility that any relationships could be driven by age-related effects. Exploratory results, uncorrected for multiple comparisons over ROIs, are detailed in the Supplementary Results and Supplementary Figures 4-5. Briefly, exploratory associations were found between greater eye looking and lower activations in the dSFG, greater mouth-looking and greater activations in the dSFG and rIFG, greater eye-looking and lower rSTG activations to angry faces, and greater mouth-looking and lTPJ activations to happy faces (Supplementary Results and Supplementary Figures 4-5). None of these effects survived FDR-correction for multiplecomparisons, and no other significant effects were found.

## Discussion

We observed significant frontal and temporal fNIRS activation to emotional faces at both 5 and 7-months of age in a paradigm that systematically varied facial identity within each facial expression block. Activations were observed in areas broadly consistent with previous adult fMRI studies, including temporal areas (Haxby et al., 2001), IFG (Sabatinelli et al., 2011), TPJ, and medial prefrontal SFG (Etkin, Egner, & Kalisch, 2011). Specifically, we found broad bilateral temporal and frontal responses to happy faces, in line with previous infant studies in which facial identity remained unvarying within each block (Grossmann et al., 2018; Krol et al., 2019; Minagawa-Kawai et al., 2009). Exploratory analyses of the current data revealed greater frontal (dSFG) activation to happy versus fearful faces, robust to changes in facial identity; this finding did not survive correction for multiple comparisons. At least one fMRI study in adults (Zhang et al., 2016) and another fNIRS study in infants (Grossmann et al., 2018) also do not report differential activations to different types of facial emotions over temporal cortices. Interestingly, and contrary to previous reports in behavioral and ERP paradigms (Leppänen & Nelson, 2009; Xie et al., 2018), no differences between 5- and 7-month-olds were found. Because ERP and fNIRS paradigm differ in paradigm design, response temporality, and underlying neural sources, some or all of these differences may account for the differences in findings. That is, 5- and 7-month-olds may differ in their fast, neurophysiological responses to isolated facial expression stimuli, but not in their slow-building hemodynamic responses to block of facial expression stimuli of varying identities. Alternatively, neural sources accounting for differential neurophysiological responses depending on age or facial expression condition as captured by ERPs may simply not be accessible by fNIRS. For example, Xie et al (2018) estimated source activity from ERP responses to facial expressions of emotion (happy, fear, anger) in 5-, 7-, and 12-month-old infants: differential responses based on emotion or age were most evident in regions of interest corresponding to the occipital face area, posterior cingulate, and fusiform face area, neither of which were directly accessible by the fNIRS apparatus used in the current study. Xie et al (2018) also reported a lack of differential responses to different types of emotional facial expressions in the superior temporal region of interests, which is consistent with the current fNIRS findings.

Behavioral studies have demonstrated that the ability to extract the emotional expression of faces across identities emerges later in development than the ability to extract the emotional expression of a single model (Bayet & Nelson, 2019). The broad neural activations to happy expressions that are reported here, occurring as face identity varied within each block, may be interpreted as further evidence that infants at this age can extract happy expressions from faces of different identities. Variations of facial identity within each block likely explain the lack of differential responses to types of emotional faces over temporal areas, in contrast to Nakato et al. (2011), and over the right IFG, in contrast to Krol et al (2019).

Infants looked marginally longer to the eyes of fearful faces, and more to the mouth of angry faces, regardless of age. These differences may reflect special salience of fearful eyes (Adolphs et al., 2005; Dadds, El Masry, Wimalaweera, & Guastella, 2008; Eisenbarth & Alpers, 2011). More looking to the mouth for angry faces has not been consistently reported in adults (Beaudry, Roy-Charland, Perron, Cormier, & Tapp, 2014). Critically, all expressions presented in this study had open “toothy” mouths, suggesting that the low-level perception of “toothiness” (Caron, Caron, & Myers, 1985) alone cannot account for this finding. We did not observe condition differences in overall looking time or heart rate responses, suggesting that emotional category did not impact arousal or attentiveness. The current block design, and variations of facial identity within each block, likely explain the differences in findings compared to earlier findings (Peltola et al., 2013; Peltola, Leppänen, & Hietanen, 2011). The current negative results are aligned with the notion that young infants may not necessarily extract or infer affective meaning from the emotional facial expressions that they can perceptually differentiate (Ruba & Repacholi, 2019).

Exploratory relationships were found between looking behavior and brain responses in the dSFG, rIFG, rSTG, and lTPJ. As none survived correction for multiple comparisons over ROIs, these findings will be discussed only briefly. Less looking to the eyes and more looking to the mouth was related to greater dSFG activations across all emotion conditions. The eye region is particularly relevant for decoding facial expressions (Eisenbarth & Alpers, 2011), and prefrontal deactivation is associated with visual attention in children (Fekete, Beacher, Cha, Rubin, & Mujica-Parodi, 2014), adults (Lachaux et al., 2008), and infants (Xu et al., 2017). Thus, a decrease in dSFG activations with more eye looking may reflect increased attentional engagement. An exploratory relationship between rIFG activation and greater mouth-looking was found, in line with another study reporting a positive correlation between mouth-looking and IFG activation in infants during speech perception (Altvater-Mackensen & Grossmann, 2016). Increased mouth-looking to happy faces was also associated with increased lTPJ activation, consistent with the diagnostic value of the mouth-region in decoding happy faces (Smith et al., 2005) and the sensitivity of the infant lTPJ to social stimuli (McDonald & Perdue, 2018). Greater eye-looking to angry faces was associated with decreased activation in the rSTG, perhaps reflecting the failure to adequately detect and process the angry mouth.

Our correlational approach cannot determine if differences in brain activations were causally driving differences in looking behavior, or vice-versa, and other factors might mediate the associations reported. The spatial resolution of the fNIRS apparatus used in the current study did not allow us to examine fine-grained, multivariate patterns of response that may distinguish between facial emotions in infants as they do in adults (Peelen, Atkinson, & Vuilleumier, 2010; Skerry & Saxe, 2014). Cross-sectional designs are inherently limited in their ability to detect developmental changes. Future work including high-density fNIRS in a longitudinal prospective design may provide a more complete picture of emotional face processing in infancy. While all blocks were presented while infants were looking to the screen, there is still a possibility that signals in the baseline period may contain responses to uncontrolled stimuli in the testing environment. This is a common concern with infant studies that may be alleviated in future work by attempting to enforce infant looking during a longer pre-stimulus period.

In summary, we simultaneously recorded fNIRS neural activation, eye-tracking, and heart rate responses to happy, fearful, and angry facial expressions of faces of varying identities in 5- and 7-month-old infants. Temporo-parietal and frontal activations to happy and angry faces and temporoparietal activations to fear faces were observed; activations were particularly evident in response to happy faces. Infants looked longer to the mouth region of angry than happy or fearful faces, and no differences in behavior or neural activations were observed between 5- and 7-month-olds.

## Acknowledgements

We thank the infants and their families for their participation. This work was supported by the National Institutes of Health [NIMH #MH078829 awarded to CAN, NICHD #HD018942 to JER] and a Fulbright grant awarded to HFB. Assistance with data collection was provided by Lindsay Bowman, Dana Bullister, Anna Fasman, Sarah McCormick, Lina Montoya, Ross Vanderwert, and Anna Zhou. Assistance with eye-tracking data processing was provided by Helga Miguel. Statistical advice was provided by Kush Kapur through the Harvard Catalyst program.

## Acknowledgements

The authors declare no competing interests.

