## Supplementary Materials for "Neural responses to happy, fearful and angry faces of varying identities in 5- and 7-month-old infants"

### Supplemental Results

*Exploratory fNIRS results*

Differences in oxyHb responses between ages and conditions were tested using a repeated-measures ANOVA at each ROI. No significant interaction terms were found between age and condition, so the interaction term was removed from the models. A significant effect of condition was seen in the dSFG ($F(2, 260)=4.30$, $p=.019$), uncorrected for multiple comparisons over ROIs. Post-hoc paired *t*-tests revealed greater activation to happy as opposed to fearful faces ($t(87)=3.24$, $p=0.002$, Supplementary Figure 3), but no difference between happy and angry faces ($t(87)=1.86$, $p=.066$) or between fearful and angry faces ($t(87)=0.61$, $p=.54$). As reported in the main manuscript, however, this effect did not survive FDR-level correction for multiple comparisons over ROIs. No main effect of age was seen in any ROI.

*Exploratory combined fNIRS and eye-tracking results*

Differences in oxyHb responses based on looking time were tested using repeated-measures ANOVAs in each ROI. The model was estimated separately for eye- and mouth-looking. Fixed effects were condition, looking time to the eyes or mouth, and the interaction between condition and looking time. Exploratory results, uncorrected for multiples comparisons over ROIs, are summarized in **Supplementary Figure 4** and detailed below. None survived FDR correction for multiple comparisons.
For eye-looking, a significant interaction between condition and looking time was seen on brain activation in the rSTG ($F(2, 167)=4.04$, $p=.019$). Post-hoc regressions for each condition separately showed a significant negative relationship between looking to the eyes and brain activity in the rSTG for angry faces ($F(1, 55)=6.26$, $p=.015$) but not for happy faces ($F(1, 55)=.007$, $p=.93$) or fearful faces ($F(1, 55)=2.04$, $p=.16$). A condition-independent relationship between looking time and brain activity was seen in the dSFG, where eye looking had a negative relationship with activation magnitude ($F(1, 185)=5.19$, $p=.024$).
For mouth-looking, a significant interaction between condition and looking time was seen in the lTPJ ($F(2, 206)=3.22$, $p=.042$). Post-hoc regressions for each condition separately showed a significant positive relationship between looking to the mouth and brain activity in the lTPJ for happy faces ($F(1, 68)=5.71$, $p=.020$, Figure 5), but not for angry faces ($F(1, 68)=.22$, $p=.64$,) or fearful faces ($F(1, 68)=.63$, $p=.43$). A condition-independent relationship between looking time to the mouth and brain activity was seen in the dSFG and rIFG, where mouth looking had a positive relationship with activation magnitude (dSFG: $F(1, 185)=6.01$, $p=.015$, rIFG: $F(1, 194)=5.66$, $p=.018$). The relationship between mouth looking and the oxyHb response is shown in **Supplementary Figure 5**.

**Supplemental Figure 1.** *Significant oxyHb activations for each emotional category and age after correcting for multiple comparisons displayed on a 7.5-month-old MRI atlas.*

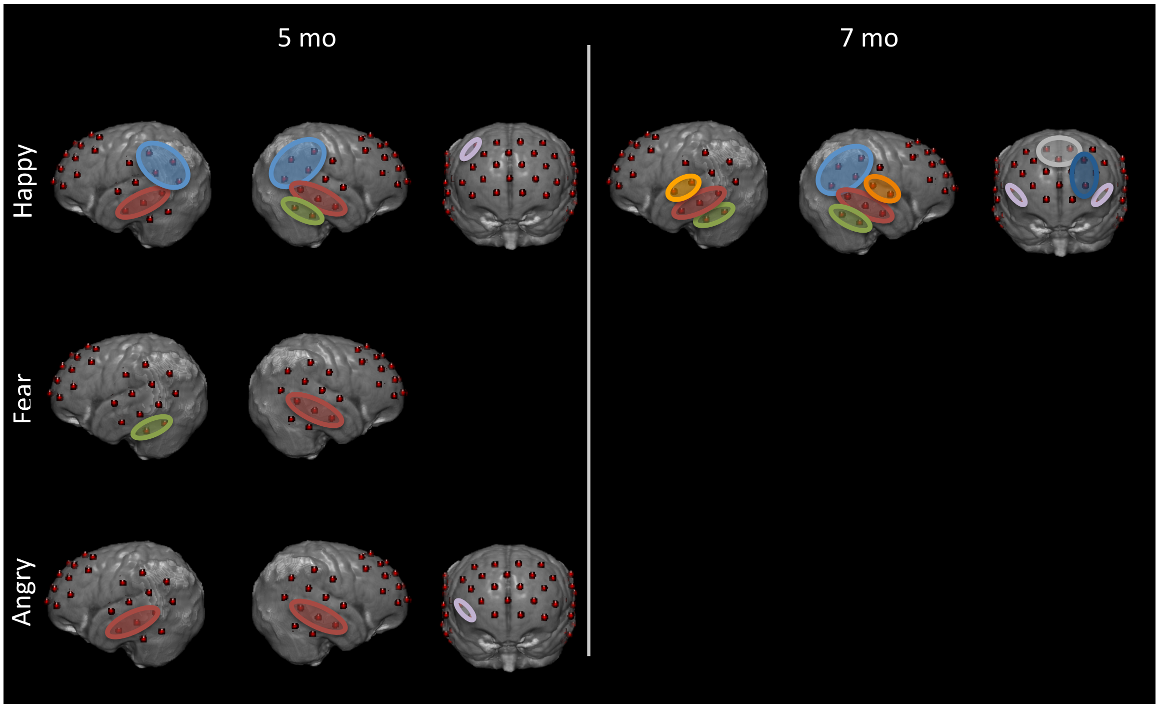


**Supplemental Figure 2.** *Example time-courses of the oxyHb response by condition and age for rTPJ, rMTG, lIFG, rMFG*

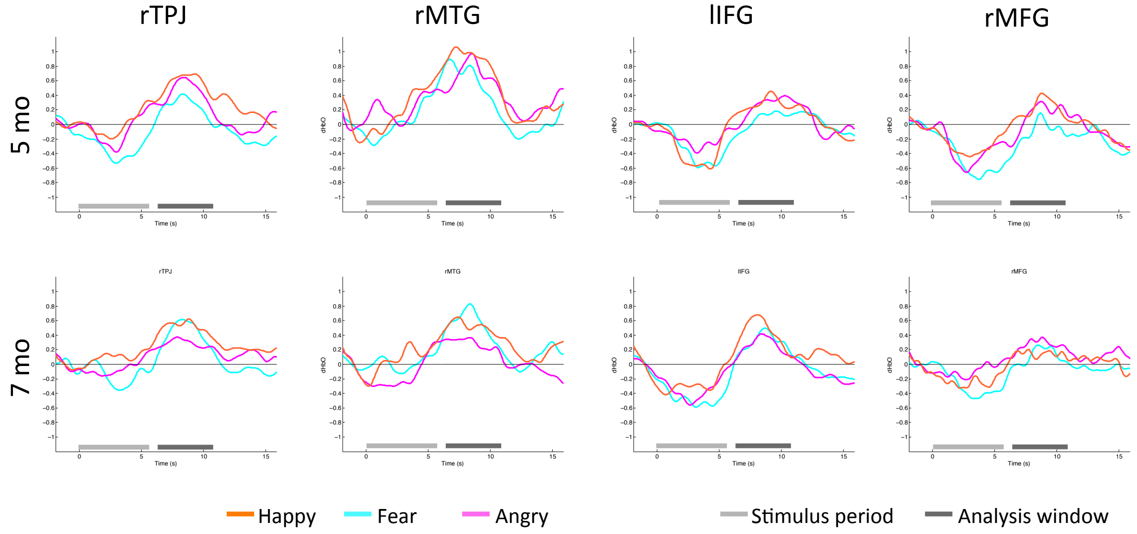

**Supplemental Figure 3.** *OxyHb response to happy and fearful faces in the dSFG ROI*

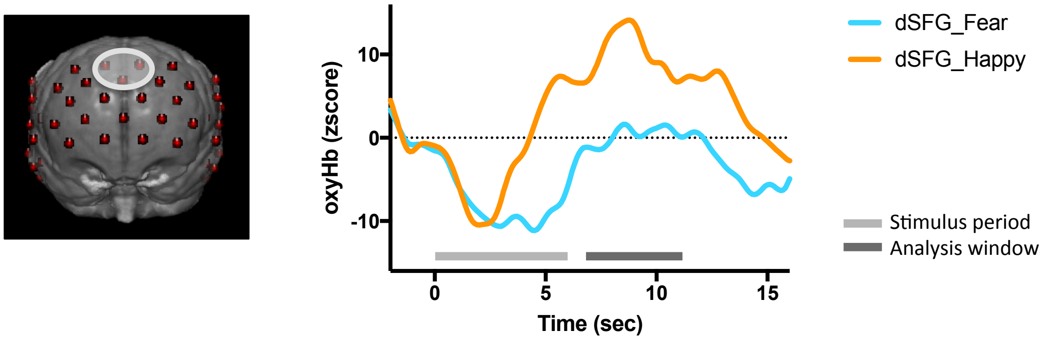


**Supplemental Figure 4.** *Significant relationships between looking time and oxyHb responses, uncorrected for multiple comparisons over ROIs.*

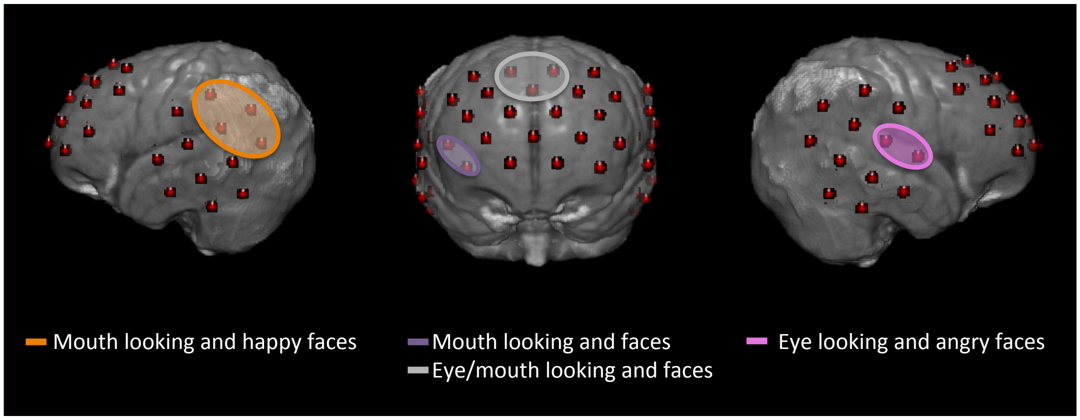


**Supplemental Figure 5**

*Left. Correlation between the oxyHb response in the lTPJ and looking time to the mouth for happy faces. Right. OxyHb response in the lTPJ for infants with long and short mouth looking (median split)*

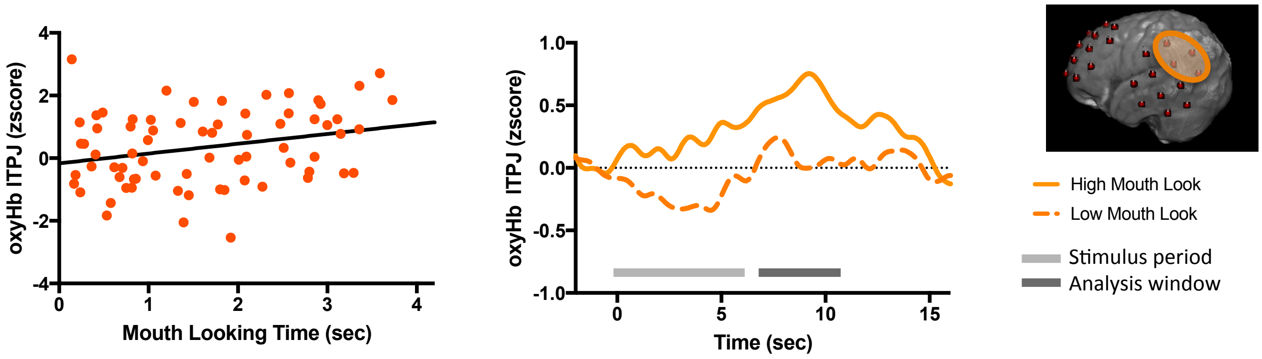

*Supplemental Table 1:* ROIs with significant activation in 5-month-old cohort

| | ROI | Uncorrected *p*-value | | | Corrected *p*-value | | |
|---|---|---|---|---|---|---|---|
| | | Happy | Fear | Angry | Happy | Fear | Angry |
| oxyHb | | | | | | | |
| Frontal | vSFG | .049 | | | | | |
| | dSFG | .041 | | | | | |
| | lMFG | | | .033 | | | |
| | rIFG | | | .0087 | | | .046 |
| | lIFG | | | .049 | | | |
| | rdMFG | .0097 | | | .026 | | |
| Temporal | rMTG | .0011 | .0013 | .0012 | .0089 | .011 | .0094 |
| | lMTG | <.001 | .027 | <.001 | .026 | | .0049 |
| | rSTG | | | | | | |
| | rTPJ | <.001 | | .041 | .0018 | | |
| | lTPJ | .0032 | | .030 | .009 | | |
| | rITG | <.001 | .017 | .046 | .026 | | |
| | lITG | | .001 | .025 | | .011 | |
| deoxyHb | | | | | | | |
| Frontal | lIFG | .032 | | | | | |
| Temporal | lMTG | .010 | .022 | | | | |
| | rTPJ | | | .017 | | | |
| | lTPJ | | | .0085 | | | |

*Supplemental Table 2:* ROIs with significant activation in the 7-month-old cohort

| | ROI | Uncorrected *p*-value | | | Corrected *p*-value | | |
|---|---|---|---|---|---|---|---|
| | | Happy | Fear | Angry | Happy | Fear | Angry |
| oxyHb | | | | | | | |
| Frontal | vSFG | | | .043 | | | |
| | dSFG | .0027 | | | .0088 | | |
| | rMFG | | .041 | | | | |
| | lMFG | .0082 | | | .017 | | |
| | rIFG | .0025 | | .042 | .0088 | | |
| | lIFG | <.001 | .0035 | .021 | .0019 | | |
| | ldMFG | | | | | | |
| | rdMFG | .039 | | | | | |
| Temporal | rMTG | <.001 | .0045 | | .0019 | | |
| | lMTG | .0019 | .036 | .013 | .0088 | | |
| | rSTG | .028 | | | .043 | | |
| | lSTG | .030 | | | | | |
| | rTPJ | .0083 | .0071 | | .017 | | |
| | lTPJ | | .024 | | | | |
| | rITG | .0086 | | | .017 | | |
| | lITG | .028 | | | .043 | | |
| deoxyHb | | | | | | | |
| Frontal | vSFG | . | | .048 | | | |
| | rMFG | .025 | | | | | |
| | lIFG | .0035 | | | | | |
| Temporal | rMTG | .048 | | .039 | | | |
| | lMTG | | .038 | .012 | | | |
| | lSTG | .032 | | | | | |
| | rlITG | .012 | | | | | |
